# An attempt of using public ambient temperature data in swine genetic evaluation for litter size traits at birth in Japan

**DOI:** 10.1101/2022.02.09.479814

**Authors:** Hitomi Hara, Shinichiro Ogawa, Chika Ohnishi, Kazuo Ishii, Yoshinobu Uemoto, Masahiro Satoh

## Abstract

To obtain the fundamental information on using ambient temperature information in developing the model for routine swine genetic evaluation in Japan, we analyzed total number born (TNB), number born alive (NBA), and number stillborn (NSB) collected at a Japanese farm, together with off-farm ambient temperature measured at a nearest Automated Meteorological Data Acquisition System station. Five repeatability animal models were exploited, considering the effects of farrowing season (model 1), farrowing month (model 2), quadratic regressions of daily maximum ambient temperature of farrowing day (model 3), season and temperature (model 4), or month and temperature (model 5). Patterns of the effects of daily maximum temperature of farrowing day estimated using model 3 was similar to those of farrowing season by model 1 and those of farrowing month by model 2. Adding the effect of daily maximum temperature of farrowing day (models 4 and 5) could explain phenotypic variability greater than only considering either of farrowing season and month (models 1 and 2). Estimated heritability was stable among the models and the rank correlation of predicted breeding values between models was >0.98 for all traits. The results indicate the possibility that using public ambient temperature can capture a large part of the phenotypic variability in litter size traits at birth caused by the seasonality in Japan and do not harm, at least, the performance of genetic evaluation. This study could support the availability of public meteorological data in flexible developing operational models for future swine genetic evaluation in Japan.

## 1. INTRODUCTION

In Japan, pork production traits, including average daily gain, longissimus muscle area, and intramuscular fat content with middle to high heritabilities, have been genetically improved by selection (e.g., Suzuki *et al*. 2005; Kadowaki *et al*. 2012; Ohnishi and Satoh 2018). Now, improving sow lifetime productivity is a pressing challenge to efficient pork production, although the heritabilities of litter size traits at birth have been estimated to be low (e.g., Tomiyama *et al*. 2011; Ogawa *et al*. 2019a; Ogawa *et al*. 2020). Potentials have been assessed of several approaches, such as choosing a statistical model more appropriate to estimate breeding values for number born alive (NBA) in terms of the parity order of dam (Ogawa *et al*. 2019b; Konta *et al*. 2020), exploring a preferable trait to assist in genetically improving NBA (Konta *et al*. 2019; Ogawa *et al*. 2019a; Ogawa *et al*. 2020), and investigating the possibility of genetically improving sow longevity (Ogawa *et al*. 2021a). A possible different approach is to perform a large-scale genetic evaluation across farms because this might predict breeding values that have higher accuracies by using more phenotypic information obtained from relatives reared on different farms and that can be directly compared between individuals on different farms. Therefore, it is important to provide an operational model suitable for a large-scale routine genetic evaluation by simultaneously using data collected from around Japan.

Japan is an island country that has four distinct seasons with a climate ranging from subarctic in the north to subtropical in the south, and the conditions are different between the Pacific side and the Sea of Japan side (https://www.data.jma.go.jp/gmd/cpd/longfcst/en/tourist.html). Japanese pig farms are widely distributed in Japan (e.g., Koike *et al*. 2018; Ogawa *et al*. 2019c; Fujimoto *et al*. 2021).

Previous studies have reported that seasons affect the meat production and reproductive performance of pigs reared in Japan (e.g., Harada *et al*. 1992; Saito and Koketsu 2009; Kakuma 2018), and statistical models considering the effects of season at measuring phenotypic information have been widely used in the genetic evaluation of traits that can be recorded throughout the year (e.g., Tomiyama *et al*. 2011; Kadowaki *et al*. 2019; Ogawa *et al*. 2019a; Ogawa *et al*. 2020). The scale of national swine genetic evaluation is getting larger (http://www.nlbc.go.jp/kachikukairyo/iden/buta/chiikinai.html), and the current evaluation uses operational models including the fixed cross-classified effects of the combination of region and season at measuring (http://www.nlbc.go.jp/kachikukairyo/iden/buta/hyokaho.html). However, these models could not explain the variation by the time of year within each season.

Effects of the time of year on pigs’ trait expression could be associated with ambient climate conditions, including thermal environment. Using information on thermal environment, such as ambient temperature, might explain not only the variation across seasons but also that within each season. However, full details of thermal environment for farm animals are rarely available. On the other hand, the utility of public meteorological data has been studied worldwide. For instance, by using off-farm temperature data measured at weather stations, Zumbach *et al*. (2008) analyzed carcass weight of pigs raised on 2 farms in North Carolina, USA; Lewis and Bunter (2011) analyzed several production traits of gilts and litter traits of sows on a farm in Australia; Tummaruk (2012) analyzed age at first observed estrus in gilts in 4 commercial herds in Thailand; Wegner *et al*. (2014) and Wegner *et al*. (2016) analyzed the numbers of total born, liveborn piglets, stillborn piglets, and weaned piglets of sows on several farms in Germany; and Mellado *et al*. (2018) analyzed the numbers of live pigs, stillborn pigs, and mummified pigs of gilts and sows on a single farm in central west Mexico. Freitas *et al*. (2006) reported that correlations between the temperature humidity index (THI) according to on-farm temperature and humidity records on a Holstein farm in Tifton, Georgia, USA, and THI according to records measured at several weather stations (up to 300 km apart) were > 0.92. Following this, Bloemhof *et al*. (2013) showed the utility of daily maximum temperature from weather stations as a heat stress indicator for farrowing rate and total number born on 16 farms in Spain and Portugal. Using public meteorological data, other studies reported the results of epidemiological investigations of total number born, weaning-to-first-mating interval, adjusted 21-day litter weight, peripartum pig deaths, and farrowing rates of gilts and sows (mainly crossbred) reared in Japanese commercial farms, in terms of herd management (e.g., Iida and Koketsu 2013; Iida and Koketsu 2014a; Iida and Koketsu 2014b; Sasaki *et al*. 2018). Previous studies used information on daily maximum temperature to investigate the effect of ambient temperature on reproductive traits of pig (e.g., Lewis and Bunter 2011; Bloemhof *et al*. 2013; Iida and Koketsu 2013; Iida and Koketsu 2014a; Iida and Koketsu 2014b; Sasaki *et al*. 2018).

No study has been investigated in detail about the performance of genetic evaluation of Japanese purebred pig population using public ambient temperature information. This is a challenging task, and as a first step, it seems reasonable to start from assessing the performance of using public meteorological data in swine genetic evaluation by analyzing data from a single farm, owing to secure the interpretability of the results. Total number born (TNB), NBA, and number stillborn (NSB) at birth are the traits having been repeatedly investigated for foreign pig populations (e.g., Lewis and Bunter 2011; Bloemhof *et al*. 2013; Wegner *et al*. 2014; Wegner *et al*. 2016), this could give more opportunities to make a meaningful discussion. This study, aiming at obtaining fundamental information for establishing a more efficient swine breeding scheme in Japan, compared on-farm temperature data measured on a single swine farm and off-farm data acquired at the nearest Automated Meteorological Data Acquisition System (AMeDAS) station (https://www.jma.go.jp/jma/en/Activities/amedas/amedas.html), and estimated genetic parameters of TNB, NBA, and NSB and predicted breeding values for these traits using the public temperature data.

## 2. MATERIALS AND METHODS

### 2.1 Ethics statement

All procedures involving animals were performed in accordance with the National Livestock Breeding Center’s guidelines for the care and use of laboratory animals.

### 2.2 Phenotypic information and pedigree data

We used 1,161 records for TNB, NBA, and NSB of 437 purebred Duroc dams obtained between 24 April 2010 and 8 August 2017 at the National Livestock Breeding Center’s Miyazaki Station (31°56′ N, 130°56′ E, 462 m a.s.l.) in Miyazaki Prefecture (Ogawa *et al*. 2020; Yazaki *et al*. 2020; Ogawa *et al*. 2021). Miyazaki Prefecture is located along the southeastern coast of the island of Kyusyu in Japan and has a subtropical climate. Artificial insemination was used for all service events. NBA was determined by the next day of the farrowing and included the number of piglets dead when checking to determine NBA but seemed to be alive at farrowing. The number of mummified piglets was not included in NSB. TNB was calculated as the sum of NBA and NSB. The pedigree data included information on 11,631 individuals. Basic statistics from the phenotypic records for the studied traits are listed in Table 1.

**TABLE 1.**
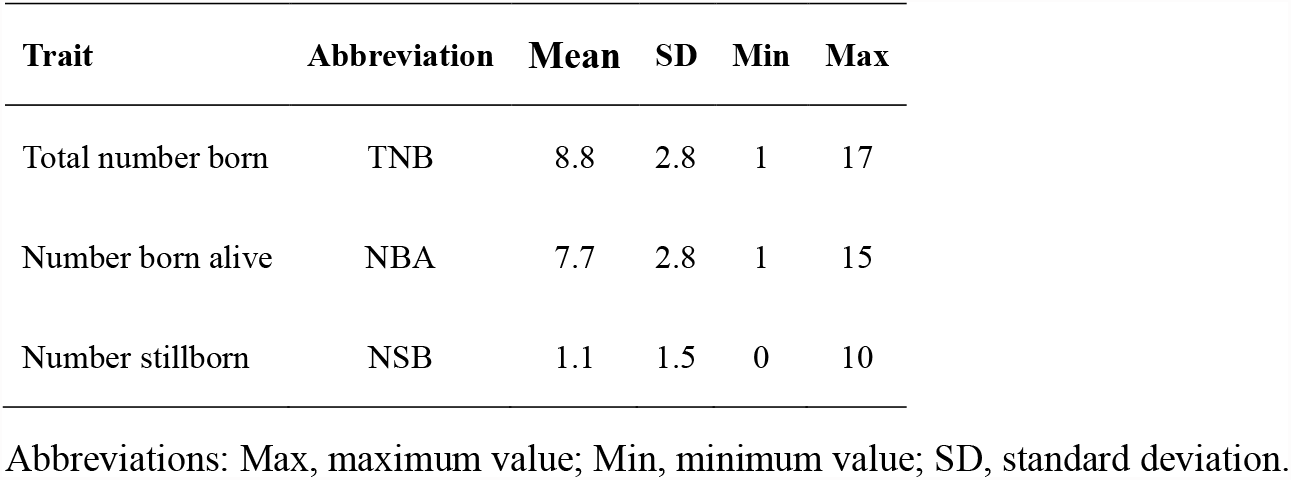
Descriptive statistics of phenotypic measurements of the studied traits

### 2.3 On- and off-farm ambient temperature data

As on-farm ambient temperature data, temperatures within two sire barns and two growing barns recorded at intervals of 5 or 10 minutes from 4 October 2016 to 24 August 2017 were available. As off-farm ambient temperature data, the daily maximum temperatures acquired by the Kobayashi AMeDAS station (32°00′ N, 130°57′ E, 276 m a.s.l.), about 6 km from the farm, from 28 December 2009 to 24 August 2017 were obtained from the Japan Meteorological Agency’s homepage (https://www.data.jma.go.jp/gmd/risk/obsdl/).

### 2.4 Numerical analyses

The following single-trait linear animal model was used:

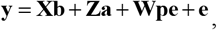

where **y** is the vector of phenotypic records; **b** is the vector of macro-environment effects (fixed effect); **a** is the vector of breeding values (random effect); **pe** is the vector of permanent environment effects (random effect); **e** is the vector of random errors (random effect); and **X, W**, and **Z** are the design matrices relating **y** to **b, a**, and **pe**, respectively. Macro-environment effects included farrowing year (2010 to 2017), parity of dam (1st to 8th), mating sire breed (Duroc, Large White), and the time of year as one of farrowing season (spring [March–May], summer [June–August], autumn [September–November], winter [December–February]; model 1), farrowing month (January to December; model 2), daily maximum temperature of farrowing day at Kobayashi AMeDAS station (quadratic regression; model 3), farrowing season and daily maximum temperature of farrowing day (model 4), or farrowing month and daily maximum temperature of farrowing day (model 5). The mean and variance–covariance of the random effects were as follows:

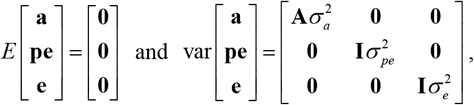

where 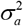 is the additive genetic variance; 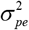 is the permanent environmental variance; 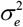 is the error variance; **A** is the additive genetic relationship matrix; and **I** is the identity matrix. Variance components were estimated in AIREMLF90 program (Misztal *et al*. 2002). Standard errors of the estimated heritability and repeatability were calculated using the elements of the inverse of the average-information matrix at convergence (Klei and Tsuruta 2008). Default setting was used as convergence criteria in the iteration procedure, that is, an iteration was stopped when at least one of the following conditions was first satisfied:

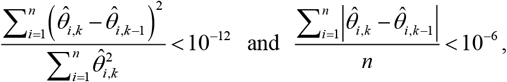

where 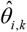 is the estimated value of parameter *i* in iteration *k* and *n* is the number of parameters to be estimated.

## 3. RESULTS AND DISCUSSION

### 3.1 Comparing on- and off-farm temperature data

Fig. 1 illustrates the relationship of on- and off-farm daily maximum temperatures. Values of Pearson’s correlation coefficients between the temperatures were very high, ranging from 0.92 to 0.96, and therefore it is reasonable to assume that off-farm daily maximum temperatures corresponded 1:1 with on-farm daily maximum temperatures in this study. The reason for such high correlations might be that the distance between the farm and the AMeDAS station is only 6 km, with no obvious geographical barrier between the farm and station (Freitas *et al*. 2006). Sasaki *et al*. (2018) used meteorological data measured at weather stations nearest each of 25 swine farms and found that the coefficient of determination between on- and off-farm temperatures was high (0.85). Our coefficients of determination ranged from 0.84 to 0.93, equal to or higher than that found by Sasaki *et al*. (2018).

**FIGURE 1.**
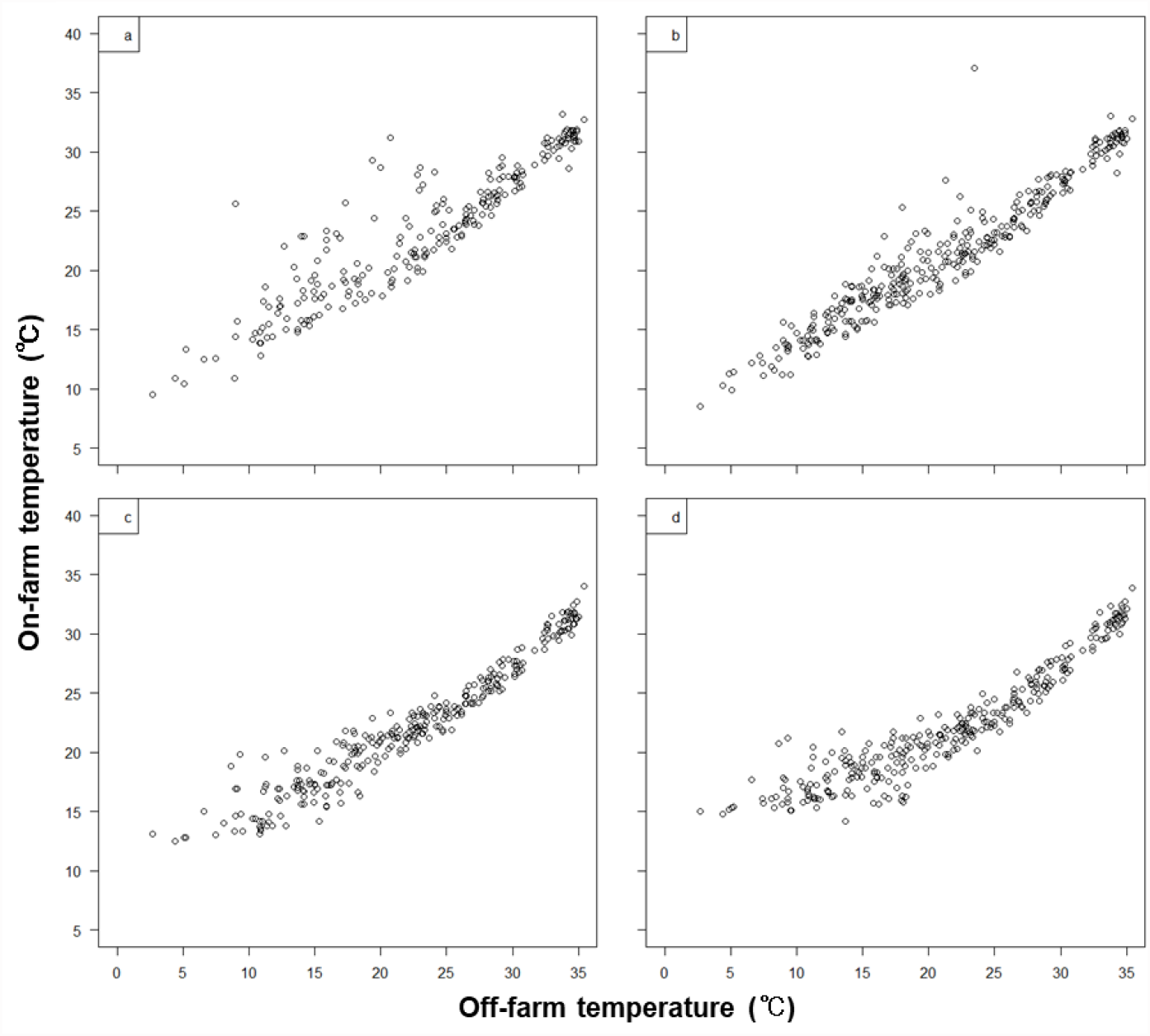
Scatter plots for on- and off-farm daily maximum temperatures. (a, b) Boar barns; (c, d) growing barns.

Nevertheless, the ranges of the values of on-farm temperatures in all barns were narrower than that of off-farm temperatures, and the distributions were slightly different between boar barns and growing barns (Fig. 1). These differences could be due to factors controlling on-farm environmental conditions. Apparent abnormal values of on-farm temperatures might reflect the effect of installation site or a flaw in the measuring devices.

On-farm temperatures were collected from October 2016, whereas most farrowing records were obtained before that date. Therefore, we used off-farm temperature data measured at Kobayashi, supposing that the 1:1 correspondence between on- and off-farm daily maximum temperatures also held before October 2016. Fig. 2 shows the relationships of off-farm daily maximum temperature of farrowing day with farrowing seasons and months. Values of the temperatures varied not only across the seasons and months in average but also within each season and month. This fact reflects the potential for capturing the phenotypic variation of traits within each season and month by using ambient temperature data. Previous study reported that the thermoneutral zone of sow was from 18 to 20°C (e.g., Curtis 1983; Peltoniemi *et al*. 1999; Bloemhof *et al*. 2013). Therefore, it could be expected that seasons and months with average values of temperature deviating from sow’ thermoneutral zone, such as summer and winter seasons and months within these seasons, might affect their phenotypic performance in this study.

**FIGURE 2.**
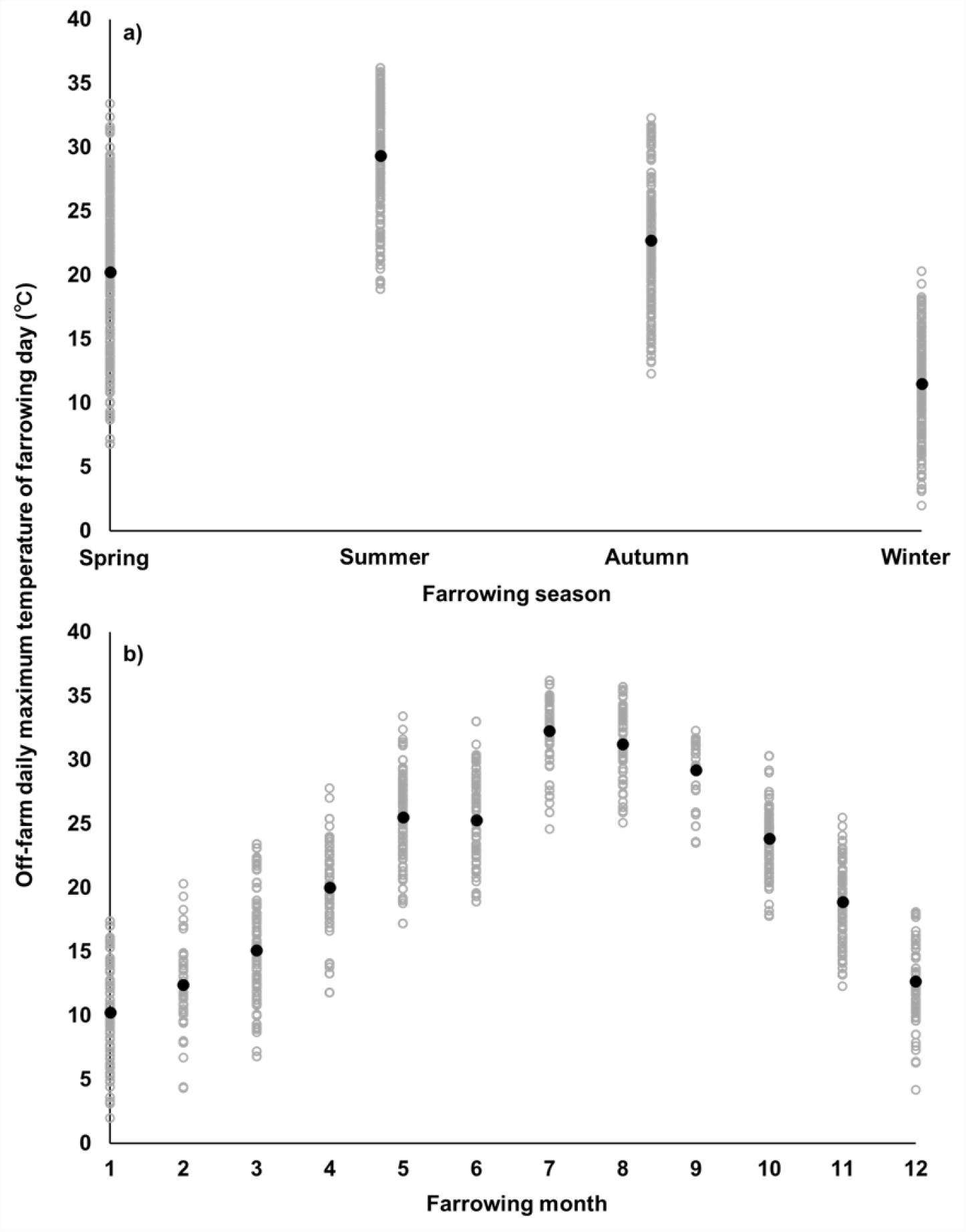
Relationships between off-farm daily maximum temperature of farrowing day and farrowing seasons and months. Black dots show the average values within each level of seasons and months.

### 3.2 Effects of farrowing season, farrowing month, and daily maximum ambient temperature of farrowing day

Table 2 summarizes the results about the estimated effects of farrowing season, farrowing month, and off-farm daily maximum temperature of farrowing day. Fig. 3 illustrates the changes in the effects of off-farm daily maximum temperature of farrowing day estimated using model 3. Fig. 4 shows the relationships of the effect of off-farm daily maximum temperature of farrowing day estimated using model 3 with corresponding farrowing seasons and months. Proportions of variances of phenotypic records explained by the estimated effects of farrowing season, farrowing month, and off-farm daily maximum temperature of farrowing day, as well as Pearson’s correlation coefficients between the values explained by those effects, are listed in Table 3. Fig. 5 shows the relationship between mating and farrowing dates and that between off-farm daily maximum temperatures of mating and farrowing days.

**TABLE 2.** Estimated values of effects of farrowing season, farrowing month, and off-farm daily maximum temperature of farrowing day

| Factor | Level | Total number born |  |  |  |  | Number born alive |  |  |  |  | Number stillborn |  |  |  |  |
| --- | --- | --- | --- | --- | --- | --- | --- | --- | --- | --- | --- | --- | --- | --- | --- | --- |
|  |  | Model |  |  |  |  | Model |  |  |  |  | Model |  |  |  |  |
|  |  | 1 | 2 | 3 | 4 | 5 | 1 | 2 | 3 | 4 | 5 | 1 | 2 | 3 | 4 | 5 |
| Season | Spring | 0.91 | - | - | 0.97 | - | 1.37 | - | - | 1.20 | - | -0.45 | - | - | -0.22 | - |
|  | Summer | 0.87 | - | - | 0.97 | - | 1.12 | - | - | 0.95 | - | -0.23 | - | - | 0.05 | - |
|  | Autumn | 0.62 | - | - | 0.69 | - | 1.03 | - | - | 0.84 | - | -0.43 | - | - | -0.16 | - |
|  | Winter | 0.00 | - | - | 0.00 | - | 0.00 | - | - | 0.00 | - | 0.00 | - | - | 0.00 | - |
| Month | January | - | 0.00 | - | - | 0.00 | - | 0.00 | - | - | 0.00 | - | 0.00 | - | - | 0.00 |
|  | February | - | 0.58 | - | - | 0.69 | - | 0.24 | - | - | 0.20 | - | 0.34 | - | - | 0.48 |
|  | March | - | 1.07 | - | - | 1.26 | - | 1.52 | - | - | 1.45 | - | -0.43 | - | - | -0.17 |
|  | April | - | 1.14 | - | - | 1.47 | - | 1.60 | - | - | 1.49 | - | -0.49 | - | - | -0.05 |
|  | May | - | 1.32 | - | - | 1.72 | - | 1.72 | - | - | 1.58 | - | -0.39 | - | - | 0.14 |
|  | June | - | 1.40 | - | - | 1.80 | - | 1.71 | - | - | 1.57 | - | -0.29 | - | - | 0.24 |
|  | July | - | 1.34 | - | - | 1.71 | - | 1.63 | - | - | 1.49 | - | -0.26 | - | - | 0.24 |
|  | August | - | 0.76 | - | - | 1.14 | - | 0.82 | - | - | 0.68 | - | -0.05 | - | - | 0.46 |
|  | September | - | 1.21 | - | - | 1.61 | - | 1.75 | - | - | 1.61 | - | -0.53 | - | - | 0.01 |
|  | October | - | 0.62 | - | - | 1.00 | - | 1.09 | - | - | 0.96 | - | -0.50 | - | - | 0.01 |
|  | November | - | 1.03 | - | - | 1.34 | - | 1.28 | - | - | 1.18 | - | -0.28 | - | - | 0.13 |
|  | December | - | 0.46 | - | - | 0.58 | - | 0.62 | - | - | 0.58 | - | -0.17 | - | - | -0.02 |
| Temperature | Linear | - | - | 0.08 | -0.02 | -0.08 | - | - | 0.19 | 0.06 | 0.02 | - | - | -0.12 | -0.08 | -0.10 |
| | Quadratic ($\times 10^2$) | - | - | -0.13 | 0.03 | 0.14 | - | - | 0.37 | 0.13 | 0.04 | - | - | 0.25 | 0.16 | 0.18 |

*Note:* Values were adjusted so that the estimated values of winter were 0 for Models 1 and 4 and those of January were 0 for Models 2 and 5.

**TABLE 3.** Proportions of variances of phenotypic records explained by the estimated effects of farrowing season, farrowing month, and off-farm maximum temperature of farrowing day (diagonal) and Pearson’s correlation coefficients between the values explained by those effects (below diagonal)

| Model | Total number born |  |  |  |  | Number born alive |  |  |  |  | Number stillborn |  |  |  |  |
| --- | --- | --- | --- | --- | --- | --- | --- | --- | --- | --- | --- | --- | --- | --- | --- |
|  | 1 | 2 | 3 | 4 | 5 | 1 | 2 | 3 | 4 | 5 | 1 | 2 | 3 | 4 | 5 |
| 1 | <u>1.40%</u> |  |  |  |  | <u>2.87%</u> |  |  |  |  | <u>1.33%</u> |  |  |  |  |
| 2 | 0.827 | <u>2.06%</u> |  |  |  | 0.880 | <u>3.66%</u> |  |  |  | 0.842 | <u>1.94%</u> |  |  |  |
| 3 | 0.642 | 0.574 | <u>0.53%</u> |  |  | 0.619 | 0.601 | <u>1.62%</u> |  |  | 0.579 | 0.524 | <u>1.41%</u> |  |  |
| 4 | 0.977 | 0.821 | 0.769 | <u>2.31%</u> |  | 0.987 | 0.883 | 0.723 | <u>2.91%</u> |  | 0.854 | 0.743 | 0.885 | <u>1.82%</u> |  |
| 5 | 0.863 | 0.972 | 0.727 | 0.889 | <u>3.71%</u> | 0.879 | 0.999 | 0.631 | 0.889 | <u>3.66%</u> | 0.730 | 0.873 | 0.760 | 0.857 | <u>2.55%</u> |

**FIGURE 3.**
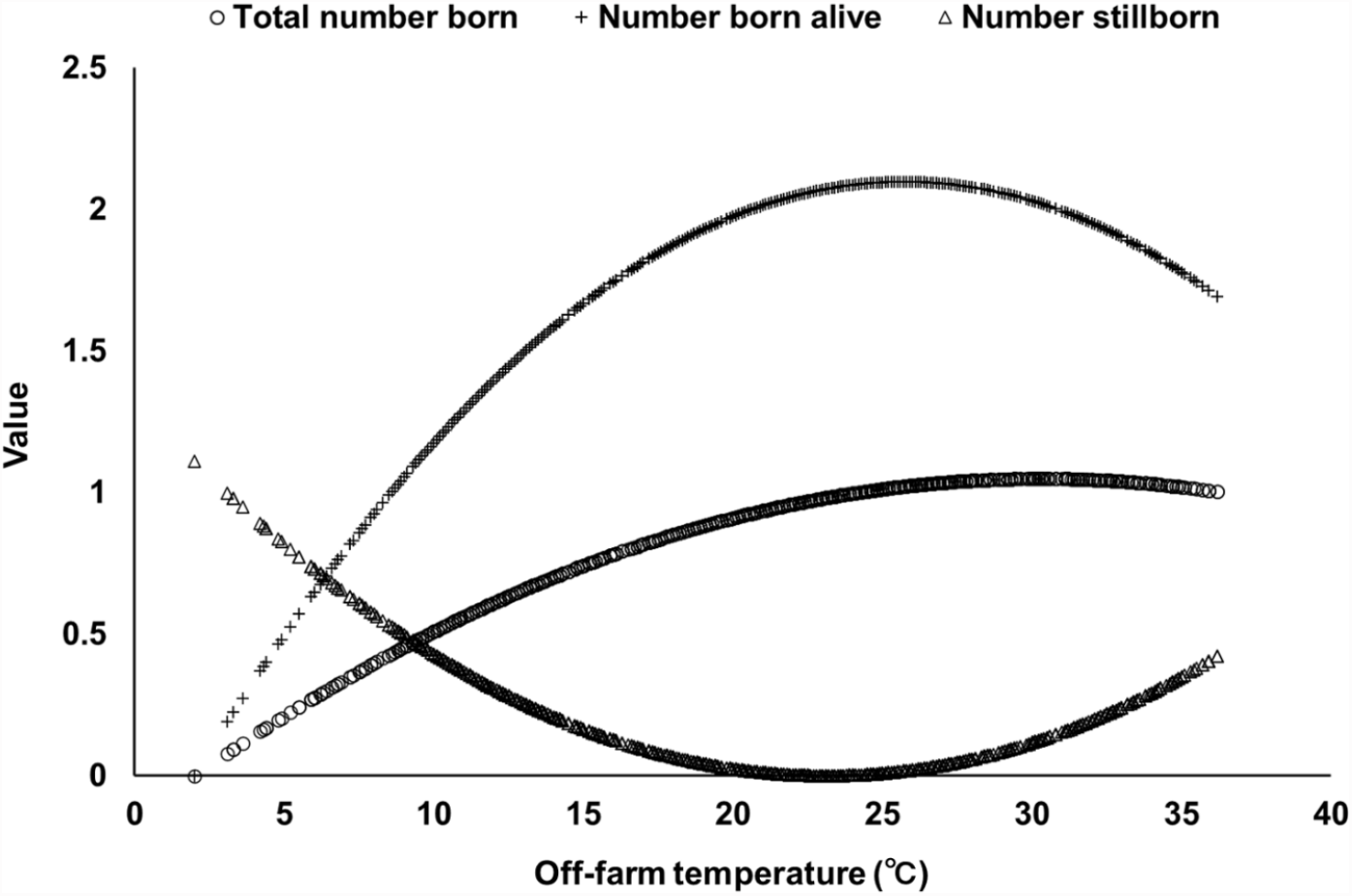
Changes in the effects of off-farm maximum temperature of farrowing day estimated using model 3. Values were adjusted so that their minimum value was equal to zero.

**FIGURE 4.**
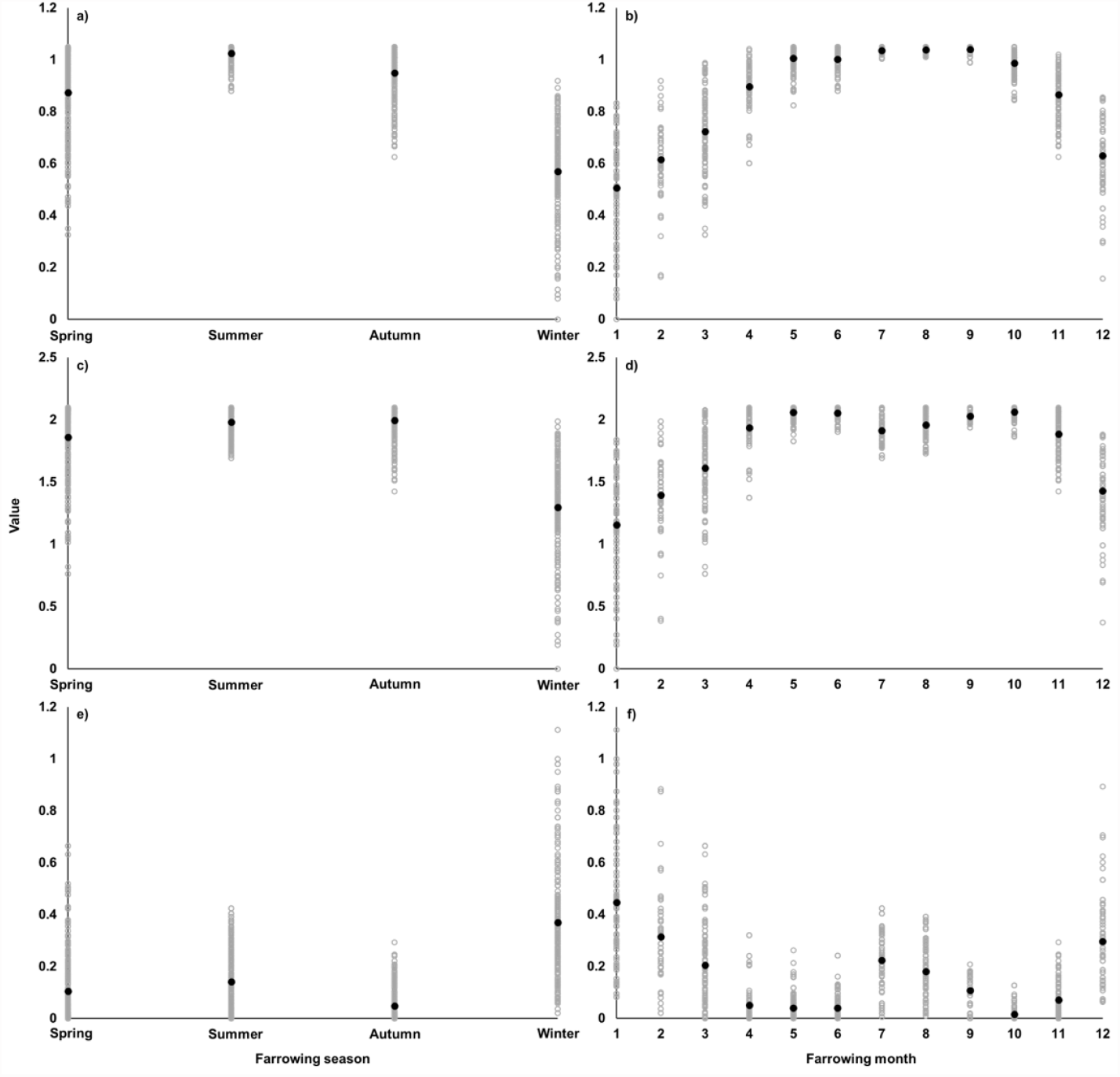
Relationships between the effect of off-farm maximum temperature of farrowing day estimated using model 3 and corresponding farrowing seasons and months for total number born (a and b), number born alive (c and d), and number stillborn (e and f). Black dots show the average values within each level of seasons and months. Values were adjusted so that their minimum value was equal to zero.

**FIGURE 5.**
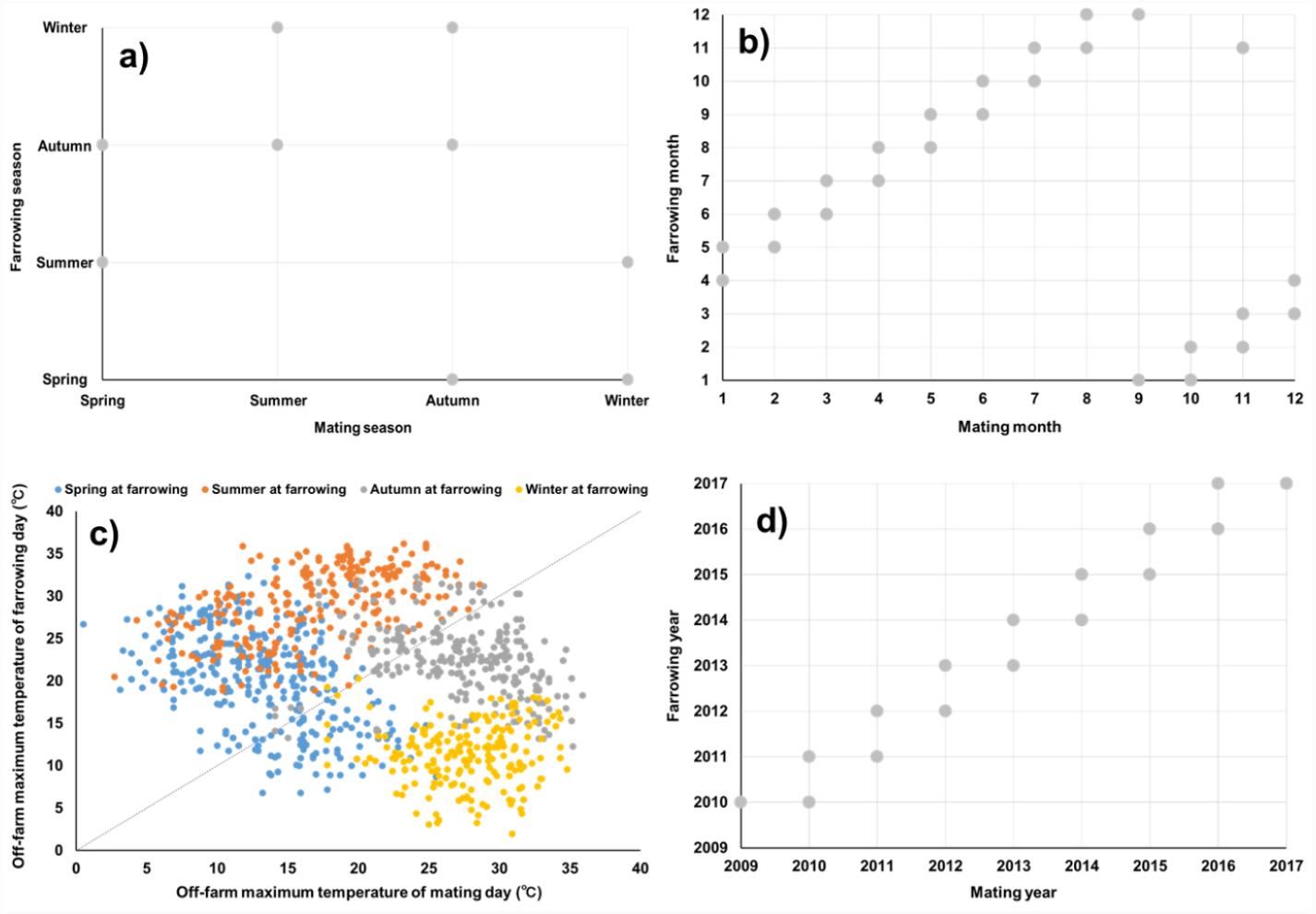
Relationship between mating and farrowing seasons (a), mating and farrowing months (b), off-farm maximum temperatures of mating and farrowing days (c), and mating and farrowing years (d) for each of farrowing records. Some records showed same month (November in autumn) as mating and farrowing months, possibly due to errors in mating dates.

Values of the effects of spring and summer at farrowing on TNB estimated using model 1 were similar to each other, that of autumn was slightly lower than those of spring and summer, and that of winter was the lowest (Table 2). TNB would be largely determined by ovulation rate, early embryonic mortality, and early fetal death (e.g., Edwards *et al*. 1968; Wildt *et al*. 1975; Nardone *et al*. 2006). Considering that the average value of gestation length in our population was 115.9 days, nearly 4 months, and that >90% of the farrowing records exhibited the gestation length ranging from 114 to 118 days (Fig. S1), most litters farrowed in winter had been artificially inseminated from August to October, hotter months in this study (Fig. 2). A heat-stressed dam before and after insemination could exhibit lower feed intake, diminished follicle stimulating hormone (FSH) and luteinizing hormone (LH) secretion, and increased body temperature and oxidative stress, these might cause lower ovarian function and higher early embryonic mortality (e.g., Flowers *et al*. 1989; Flowers and Day 1990; Quesnel *et al*. 1998; Kim *et al*. 2013). Previous studies have shown that the temperature in the period between several days prior to and after insemination had greater effects on TNB (Omtvedt *et al*. 1971; Bloemhof *et al*. 2013; Wegner *et al*. 2016).

Values of the estimated effects of spring and autumn at farrowing on NSB were similar, that of summer was slightly higher than those of spring and autumn, and that of winter was the highest (Table 2). It has been reported that heat stress in later pregnancy increased the number of stillborn piglets (e.g., Edwards *et al*. 1968; Omtvedt *et al*. 1971; Wegner *et al*. 2016). Therefore, the slightly lower value for summer at farrowing might be due to heat stress in dams. On the other hand, the lowest value for winter might be caused by cold stress in not only dam but also piglet. For example, previous studies reported the range of comfortable temperature of 18°C to 23°C for lactating sow (Yan and Yamamoto 2000; Brown-Brandl *et al*. 2001) and the minimum comfortable temperature of around 34°C to 35°C for newborn piglet (Mount 1959; Manno *et al*. 2005). Considering the physiological temperature in dam’s utero, between 38°C to 40°C, piglets encounter a colder environment immediately, this triggers the reduction of body temperature soon, known as hypothermia (Tuchscherer *et al*. 2000; Pandorfi *et al*. 2005; Malmkvist *et al*. 2006).

Values of the estimated effects for NBA were like the difference between those for TNB and NSB because TNB was the sum of NBA and NSB in this study. From what has been discussed above, one can add the effects about the time at mating, such as mating year and either of mating season, mating month, or temperature of mating day. However, care must be taken to accurate data collection about the mating and farrowing dates and a characteristic data structure about these factors (Fig. 5), due to seasonal variation in ambient temperature and less variability in gestation length. Ogawa *et al*. (2019c) reported the estimated effects of farrowing season for NBA in Landrace and Large White dams raised on multiple farms in different prefectures of a single Japanese pig breeding company. Ogawa *et al*. (2019c) showed lower estimated values for winter than for spring, although the difference between the values was 0.30 in Landrace and 0.21 in Large White, both smaller than this study. This inconsistency might be due to differences in breed, farm location, and rearing condition. Our population was reared in Miyazaki Prefecture, in southern Japan, and analyzing data obtained at different locations, such as northern Japan, might yield different effects of the time of year. Tummaruk *et al*. (2004) estimated the effects of farrowing month on NBA in Landrace and Large White populations in Thailand, showing that NBA was significantly lower in August and September than from November to June. The inconsistency might be due in part to the difference in climate conditions between countries. Bertoldo *et al*. (2012) observed that the estimated effects of season on litter size varied among studies, possibly owing to confounding factors, including parity of dam and semen characteristics. Tummaruk *et al*. (2004) and Tummaruk *et al*. (2010) reported that the effect of season on litter size was more prominent in gilts than in sows, although it is possible that their results were affected by culling for reproductive performance at earlier parities (Sasaki *et al*. 2018).

Variations in phenotypic records explained by farrowing season in model 1 and that by farrowing month in model 2 was largely in common, although the proportion on variance explained by farrowing month was greater than that explained by farrowing season for all traits (Table 3). Considering the effect of the time of year as farrowing month could explain additional variations which are not explained by farrowing season, though the statistical model becomes more complicated, that is, the number of levels increases from 4 to 12, and thus the average number of records per level is decreased. Therefore, in terms of data connectedness and reliability of the results, more careful consideration is required to interpret the results obtained using more complicated model, especially when the data structure is severely unbalanced and the data size is small. We assumed no interaction between farrowing year and farrowing season or month, and the average number of records per level decreases further when considering farrowing year-by-season and year-by-month effects.

The quadratic curve for the effect of off-farm daily maximum temperature of farrowing day estimated using model 3 was convex upward for TNB and NBA and downward for NSB (Fig. 3), although the value of the effect for TNB became stable when the temperature was >20°C. Phenotypic variation explained by the temperature in model 3 was partly in common with those explained by farrowing season in model 1 and farrowing month in model 2 (Table 3). The proportions of variances explained by the temperature were the lowest for TNB and NBA, while it was lower than that explained by farrowing month but greater than that explained by farrowing season for NSB.

Model 3 assumes that the effect of the time of year is the same when the daily maximum temperatures are the same on different days. However, sows farrowed in spring and sows farrowed in autumn might be differently affected, partly because of different responses to daylight hours, daily temperature range, and relative humidity, even if the daily maximum temperatures are the same. Therefore, we performed the analyses using models 4 and 5. For TNB and NSB, simultaneous considering the effects of the temperature and season or month (models 4 and 5) could explain more proportions of variance than considering only one of the effects of season, month, and temperature (models 1, 2, and 3) (Table 3). This indicates the availability of using temperature data to explain additional variation which could not capture by considering the effects of season and month. On the other hand, interpretating the respective values of effects of farrowing season, farrowing month, and daily maximum temperature of farrowing days estimated using models 4 and 5 were more difficult than those estimated using the other models (Table 2, Fig. S5). This could be due to confounding; that is, there are overlaps among the variations explained by farrowing season, month, and daily maximum temperature of farrowing day. Lewis and Bunter (2011) discussed the possibility of confounding when dissociating the effects of contemporary group and daily maximum temperature.

We adopted the quadratic regression of phenotypic values on temperatures, according to the results from the preliminary analysis and aiming to prevent over-fitting (Figs. S1, S2, S3, and S4). Including quadratic regression of daily maximum temperature of farrowing day lost 2 degrees of freedom, whereas including discrete effects of farrowing season and farrowing month lost 4 and 12 degrees of freedom, respectively. Furthermore, considering farrowing season and month would reduce both the average number of records per level the connectedness with other effects. On the other hand, considering daily maximum temperature as covariates in the model restricts the expression of the effect of the time of year. Therefore, modeling should be flexible in response to the structure and size of the data analyzed.

### 3.3 Genetic parameter estimation and breeding value prediction

Results of estimating genetic parameters are listed in Table 4. Values of Spearman’s rank correlation coefficients of predicted breeding values the 437 sows with their own records between the models are shown in Table 5. Values of estimated heritability and repeatability were stable across the models for all traits, mainly due to the small proportions of variance in phenotypic records explained by farrowing month, farrowing year, and daily maximum temperature of farrowing day (Table 3). Values of Spearman’s rank correlation coefficients were >0.98. These results could imply the possibility that including temperature information into the analytical model at least did not harm the performance of breeding value prediction.

**TABLE 4.** Estimated values of genetic parameters (standard errors in parentheses)

| Model | $\sigma_a^2$ | $\sigma_{pe}^2$ | $\sigma_e^2$ | $h^2$ | $rep^2$ |
| --- | --- | --- | --- | --- | --- |
| *Total number born* |  |  |  |  |  |
| 1 | 0.76 (0.43) | 0.62 (0.38) | 6.31 (0.31) | 0.10 (0.05) | 0.18 (0.03) |
| 2 | 0.88 (0.45) | 0.56 (0.39) | 6.29 (0.32) | 0.11 (0.06) | 0.19 (0.03) |
| 3 | 0.69 (0.43) | 0.67 (0.39) | 6.38 (0.32) | 0.09 (0.05) | 0.18 (0.03) |
| 4 | 0.77 (0.43) | 0.62 (0.38) | 6.32 (0.32) | 0.10 (0.05) | 0.18 (0.03) |
| 5 | 0.88 (0.45) | 0.57 (0.39) | 6.29 (0.32) | 0.11 (0.06) | 0.19 (0.04) |
| *Number born alive* |  |  |  |  |  |
| 1 | 1.20 (0.52) | 0.42 (0.40) | 6.27 (0.32) | 0.15 (0.06) | 0.20 (0.04) |
| 2 | 1.32 (0.50) | 0.30 (0.40) | 6.27 (0.31) | 0.17 (0.06) | 0.21 (0.04) |
| 3 | 1.16 (0.50) | 0.38 (0.40) | 6.40 (0.32) | 0.15 (0.06) | 0.19 (0.04) |
| 4 | 1.19 (0.50) | 0.41 (0.40) | 6.28 (0.32) | 0.15 (0.06) | 0.20 (0.04) |
| 5 | 1.33 (0.52) | 0.29 (0.40) | 6.29 (0.32) | 0.15 (0.06) | 0.20 (0.04) |
| *Number stillborn* |  |  |  |  |  |
| 1 | 0.17 (0.10) | 0.12 (0.10) | 1.90 (0.09) | 0.08 (0.05) | 0.13 (0.03) |
| 2 | 0.18 (0.10) | 0.12 (0.10) | 1.90 (0.09) | 0.08 (0.05) | 0.13 (0.03) |
| 3 | 0.17 (0.10) | 0.11 (0.10) | 1.90 (0.09) | 0.08 (0.05) | 0.13 (0.03) |
| 4 | 0.18 (0.10) | 0.11 (0.10) | 1.90 (0.09) | 0.08 (0.05) | 0.13 (0.03) |
| 5 | 0.18 (0.10) | 0.11 (0.10) | 1.89 (0.09) | 0.08 (0.05) | 0.13 (0.03) |

*Note:* $\sigma_a^2$ , additive genetic variance; $\sigma_{pe}^2$ , permanent environmental variance; $\sigma_e^2$ , error variance; $h^2$ , heritability; $rep^2$ , repeatability.

**TABLE 5.** Spearman’s rank correlations of predicted breeding values of 437 sows with their own records between different models

| Model | Total number born |  |  |  | Number born alive |  |  |  | Number stillborn |  |  |  |
| --- | --- | --- | --- | --- | --- | --- | --- | --- | --- | --- | --- | --- |
|  | 1 | 2 | 3 | 4 | 1 | 2 | 3 | 4 | 1 | 2 | 3 | 4 |
| 2 | 0.997 |  |  |  | 0.997 |  |  |  | 0.993 |  |  |  |
| 3 | 0.996 | 0.991 |  |  | 0.991 | 0.990 |  |  | 0.994 | 0.990 |  |  |
| 4 | >0.999 | 0.997 | 0.996 |  | >0.999 | 0.997 | 0.993 |  | 0.995 | 0.996 | 0.997 |  |
| 5 | 0.996 | >0.999 | 0.990 | 0.996 | 0.997 | >0.999 | 0.990 | 0.997 | 0.987 | 0.997 | 0.991 | 0.996 |

### 3.4 General discussion

With the aim of increasing the rate of genetic improvement, it is important to develop an operational model suitable for routine large-scale genetic evaluation that can handle integrated data collected around Japan. In this study, we used public meteorological data as a source of climate information to analyze phenotypic records of TNB, NBA, and NSB. This is the first study to assess the performance of using public ambient temperature data in swine genetic evaluation in Japan. We revealed that adding the effect of temperature could explain additional variations that did not explain by considering only the effect of season (Table 3). Possible reasons for our results would be that high one-to-one correspondence of off-farm temperature data with on-farm temperature (Fig. 1) and that the values of temperature varied within each season (Fig. 2). On the other hand, it should be noted that such correspondence might not be guaranteed on other farms, depending on physical and management characteristics. Therefore, it is important to investigate factors affecting the relationship between on- and off-farm ambient temperatures. Exploring a more appropriate use of temperature data, such as daily average and minimum temperatures, daily temperature range, and other factors including relative humidity, as well as choosing a different expression of the effect of ambient temperature, including linear-plateau regression and smoothing spline, might explain more phenotypic variation (Zumbach *et al*. 2008; Lewis and Bunter 2011; Bloemhof *et al*. 2013; Guy *et al*. 2017; Tiezzi *et al*. 2020). Moreover, AMeDAS relative humidity data and mesh climate data are now available as different sources of climate information. Effects of time of year, and even those of year, might vary among regions and farms. If an interaction of region and farm with time could be explained by using public meteorological data, the statistical modelling might become simpler and more objective. To tackle these challenging issues, further analysis should be performed to investigate the performance of the integrated use of different kinds of public meteorological data with a larger data set with a more complex structure.

Recently, studies have investigated the methodology for efficient breeding using information obtained from public databases; for example, genomic prediction incorporating biological information (e.g., Melzer *et al*. 2013; Ogawa *et al*. 2015; Okada *et al*. 2018) and prediction of breeding value by exploiting public meteorological data (e.g., Zumbach *et al*. 2008; Fragomeni *et al*. 2016a; Tiezzi *et al*. 2020). For the letter, it is likely that a sow’s response to high ambient temperature is somewhat heritable and that the genetic correlations of sow reproductive traits among seasons and ambient temperatures are not unity (e.g., Bloemhof *et al*. 2008; Lewis and Bunter 2011; Tiezzi *et al*. 2020). Similar results were also reported for production traits (e.g., Lewis and Bunter 2011; Fragomeni *et al*. 2016b; Usala *et al*. 2021). In pig breeding, selection has been performed to improve meat production and number of piglets weaned both in Japan (e.g., Suzuki *et al*. 2005; Tomiyama *et al*. 2011; Ohnishi and Satoh 2018) and overseas (e.g., Merks 2000; Hill 2016; Zak *et al*. 2017). Genetic improvement of productivity has brought about a large increase in total metabolic heat production (e.g., Cabezón *et al*. 2016; Johnson 2018; Johnson *et al*. 2019), reducing individual animals’ ability to cope with high ambient temperatures (e.g., Brown-Brandl *et al*. 2001; Renaudeau *et al*. 2011; Robbins *et al*. 2021). Pork production in Japan is anticipated to be affected by global warming (Takada *et al*. 2008; Sakatani 2014). Therefore, it is an urgent priority to establish new pig breeding schemes to confront global warming (e.g., Bloemhof *et al*. 2012; Schauberger *et al*. 2019; Tiezzi *et al*. 2020). In this regard, public meteorological data might offer a powerful resource, and therefore, it is important to develop a future breeding scheme to genetically improve heat tolerance of pigs in Japan (Carabaño *et al*. 2019; Mayorga *et al*. 2019; Rauw *et al*. 2020).

## DATA AVAILABILITY STATEMENT

Restrictions apply to the availability of these data, which were used under license for this study.

## CONFLICTS OF INTERESTS

The authors declare that they have no competing interests.

## ACKNOWLEDGEMENTS

This work was supported by a grant from the Ministry of Agriculture, Forestry, and Fisheries of Japan (Development of Breeding Technology for Animal Life Production).

## Notes

### Competing Interest Statement

The authors have declared no competing interest.

